# Genomic analysis of the mesophilic Thermotogae genus *Mesotoga* reveals phylogeographic structure and genomic determinants of its distinct metabolism

**DOI:** 10.1101/322537

**Authors:** Camilla L. Nesbø, Rhianna Charchuk, Stephen M. J. Pollo, Karen Budwill, Ilya V. Kublanov, Thomas H.A. Haverkamp, Julia Foght

**Author notes:** Department of Biological Sciences, CW 405 Biological Sciences Bldg., 11455 Saskatchewan Drive, University of Alberta, Edmonton, Alberta, Canada, T6G 2E9.

## Abstract

The genus *Mesotoga*, the only described mesophilic *Thermotogae* lineage, is common in mesothermic anaerobic hydrocarbon-rich environments. Besides mesophily, *Mesotoga* displays lineage-specific phenotypes, such as no or little H_2_ production and dependence on sulfur-compound reduction, which may influence its ecological role. We used comparative genomics of 18 *Mesotoga* strains (pairwise 16S rRNA identity > 99%) and a transcriptome of *M. prima* to investigate how life at moderate temperatures affects phylogeography and to interrogate the genomic features of its lineage-specific metabolism. We propose that *Mesotoga* accomplish H_2_ oxidation and thiosulfate reduction using a sulfide dehydrogenase and a hydrogenase-complex and that a pyruvate:ferredoxin oxidoreductase acquired from *Clostridia* is responsible for oxidizing acetate. Phylogenetic analysis revealed three distinct *Mesotoga* lineages (89.6-99.9% average nucleotide identity [ANI] within lineages, 79.3-87.6% ANI between lineages) having different geographic distribution patterns and high levels of intra-lineage recombination but little geneflow between lineages. Including data from metagenomes, phylogeographic patterns suggest that geographical separation historically has been more important for *Mesotoga* than hyperthermophilic *Thermotoga* and we hypothesize that distribution of *Mesotoga* is constrained by their anaerobic lifestyle. Our data also suggest that recent anthropogenic activities and environments (e.g., wastewater treatment, oil exploration) have expanded *Mesotoga* habitats and dispersal capabilities.

**Originality-Significance Statement:** This study comprises one of the first whole-genome-based phylogeographic analyses of anaerobic mesophiles, and our data suggest that such microbes are more restricted by geography than are thermophiles (and mesophilic aerobes). This is likely to be a general trait for similar anaerobic organisms – and therefore broadly relevant to and testable in other environments. Moreover, *Mesotoga* bacteria are part of the largely understudied subsurface ecosystem that has relatively recently been recognized as a new and important biosphere. Understanding the forces responsible for the distribution of organisms in the subsurface, as well as the identification of genes responsible for *Mesotoga*’s distinct metabolism, will contribute to the understanding of these communities.

## Introduction

The genus *Mesotoga* is the only characterized mesophilic lineage within the otherwise thermophilic bacterial phylum Thermotogae (Pollo *et al.*, 2015). *Mesotoga* spp. have been isolated from and detected in polluted marine sediments, low temperature oil reservoirs, and waste water treatment facilities (Nesbø *et al.*, 2010; Hania *et al.*, 2011; Nesbø *et al.*, 2012; Hania *et al.*, 2013), and are common in anaerobic methanogenic environments (Nesbø *et al.*, 2010) where they may be involved in syntrophic acetate degradation (Nobu *et al.*, 2015). The first described member of this genus, *Mesotoga prima* MesG1Ag4.2 (hereafter, *M. prima*), was isolated from a PCB-degrading enrichment culture inoculated with sediments from Baltimore Harbor, Maryland (USA) (Nesbø *et al.*, 2006; 2012). Sequencing the genomes of *M. prima* and the very closely related *M. prima* PhosAc3 (hereafter, PhosAc3) isolated in Tunisia (Hania *et al.*, 2015) revealed larger genomes than in thermophilic Thermotogae, with more genes involved in regulatory functions and interactions with the environment (Zhaxybayeva *et al.*, 2012).

Genome size in Thermotogae inversely correlates with optimum growth temperature (Zhaxybayeva *et al.*, 2012; Pollo *et al.*, 2015). However, it is unclear how growth temperature affects other aspects of genome evolution including levels of homologous recombination. Hyperthermophilic *Thermotoga* display extremely high levels of homologous recombination, which could be a side effect of the need for DNA repair at high temperatures (Nesbø *et al.*, 2015). Nesbø *et al.* (2015) also found high levels of geneflow among all *Thermotoga* spp. genomes investigated, and that populations from similar environments have exchanged more genes than geographically close populations from different environments. For instance, *Thermotoga* genomes from oil reservoirs in Japan and in the North Sea, as well as from a continental hot spring in North America, have exchanged more genes through homologous recombination than they have with genomes from geographically closer marine vents. Moreover, the phylogeographic analysis of *Thermotoga* genomes suggested that oil reservoirs were colonized from subsurface populations rather than being buried with the sediments that mature into oil reservoirs reservoirs (a corollary of the paleosterilization hypothesis; (Wilhelms *et al.*, 2001)) (Nesbø *et al.*, 2015). Comparative genomic analyses of mesophilic Thermotogae may shed light on the role of growth temperature on recombination and phylogeography.

In addition to lower optimal growth temperature (37°C - 40°C), *Mesotoga*’s core energy metabolism also differs from that of other characterized thermophilic Thermotogae. For instance, while growth of most thermophilic Thermotogae is stimulated by adding sulfur compounds to the medium (Ravot *et al.*, 1995; Boileau *et al.*, 2016), reduction of sulfur compounds appears to be essential for growth of *Mesotoga* in pure culture and they produce little or no H_2_ (Hania *et al.*, 2011; 2013; Fadhlaoui *et al.*, 2017).

Here we compare 18 *Mesotoga* genomes obtained from isolates and single cells originating from six geographically different sites, including three low temperature continental oil reservoirs, in order to elucidate genomic markers of metabolic differences and to investigate how growth temperature influences phylogeography and prevalence of recombination. We also include in our analysis *Mesotoga* sequences from publicly available metagenomes. We compare our findings from the mesophilic *Mesotoga* to the patterns previously observed in the hyperthermophilic *Thermotoga* (Nesbø *et al.*, 2015) and infer that geographic separation has had more influence on the phylogeography of *Mesotoga*, possibly due to selective pressures of dispersal of strict anaerobes through aerobic environments. Finally, we present a model that accounts for *Mesotoga*’s distinct sulfur-dependent metabolism involving a hydrogenase complex.

## Results

### Genome sequences

We generated draft genomes for eight newly isolated *Mesotoga* strains from two oil reservoirs (H and B) in Alberta Canada and one *Mesotoga* strain from a PCB-degrading enrichment culture from Baltimore Harbor, Maryland USA (Table 1). Seven partial single cell amplified genomes (SAGs) were obtained from cells sorted from produced water from an Albertan oil reservoir (PW), a naphtha-degrading enrichment culture inoculated with sediments from an Albertan oil sands tailings pond (NAPDC), and a toluene-degrading enrichment culture inoculated with sediments from a contaminated aquifer in Colorado USA (TOLDC). We also included in our analyses the draft genome of PhosAc3, previously isolated in Tunisia (Hania *et al.*, 2015) and the closed genome of *M. prima* (Zhaxybayeva *et al.*, 2012) from Baltimore Harbor.

**Table 1.**
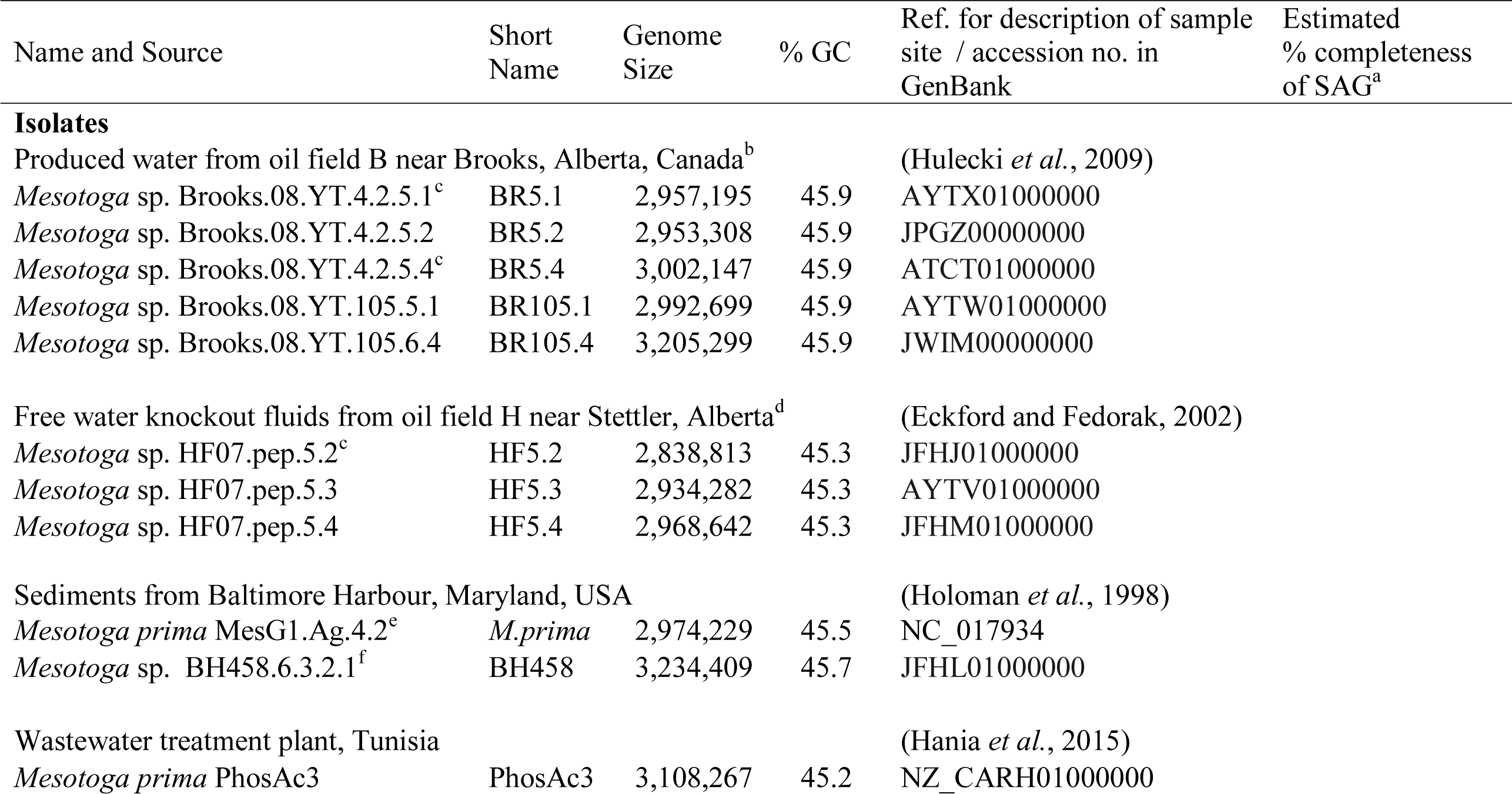
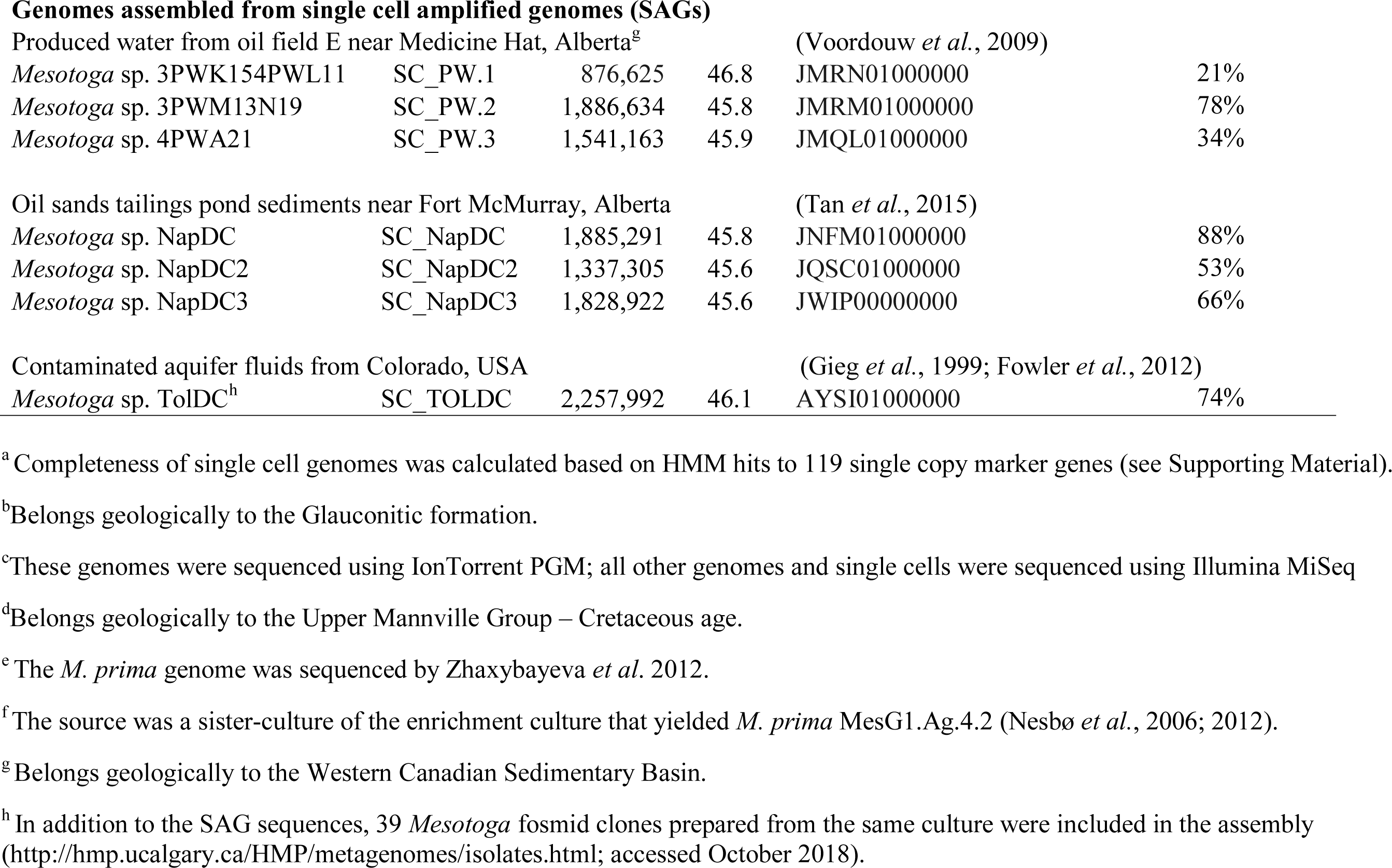
List of genomes analyzed. All genomes, except those of *Mesotoga prima* and *M. prima* PhosAc3, were sequenced as part of the current study.

| Name and Source | Short Name | Genome Size | % GC | Ref. for description of sample site / accession no. in GenBank | Estimated % completeness of SAG<sup>a</sup> |
| --- | --- | --- | --- | --- | --- |
| **Isolates** |  |  |  |  |  |
| Produced water from oil field B near Brooks, Alberta, Canada<sup>b</sup> |  |  |  | (Hulecki *et al.*, 2009) |  |
| *Mesotoga* sp. Brooks.08.YT.4.2.5.1<sup>c</sup> | BR5.1 | 2,957,195 | 45.9 | AYTX01000000 |  |
| *Mesotoga* sp. Brooks.08.YT.4.2.5.2 | BR5.2 | 2,953,308 | 45.9 | JPGZ00000000 |  |
| *Mesotoga* sp. Brooks.08.YT.4.2.5.4<sup>c</sup> | BR5.4 | 3,002,147 | 45.9 | ATCT01000000 |  |
| *Mesotoga* sp. Brooks.08.YT.105.5.1 | BR105.1 | 2,992,699 | 45.9 | AYTW01000000 |  |
| *Mesotoga* sp. Brooks.08.YT.105.6.4 | BR105.4 | 3,205,299 | 45.9 | JWIM00000000 |  |
| Free water knockout fluids from oil field H near Stettler, Alberta<sup>d</sup> |  |  |  | (Eckford and Fedorak, 2002) |  |
| *Mesotoga* sp. HF07.pep.5.2<sup>c</sup> | HF5.2 | 2,838,813 | 45.3 | JFHJ01000000 |  |
| *Mesotoga* sp. HF07.pep.5.3 | HF5.3 | 2,934,282 | 45.3 | AYTV01000000 |  |
| *Mesotoga* sp. HF07.pep.5.4 | HF5.4 | 2,968,642 | 45.3 | JFHM01000000 |  |
| Sediments from Baltimore Harbour, Maryland, USA |  |  |  | (Holoman *et al.*, 1998) |  |
| *Mesotoga prima* MesG1.Ag.4.2<sup>e</sup> | *M. prima* | 2,974,229 | 45.5 | NC_017934 |  |
| *Mesotoga* sp. BH458.6.3.2.1<sup>f</sup> | BH458 | 3,234,409 | 45.7 | JFHL01000000 |  |
| Wastewater treatment plant, Tunisia |  |  |  | (Hania *et al.*, 2015) |  |
| *Mesotoga prima* PhosAc3 | PhosAc3 | 3,108,267 | 45.2 | NZ_CARH01000000 |  |
| Genomes assembled from single cell amplified genomes (SAGs) |  |  |  |  |  |
| Produced water from oil field E near Medicine Hat, Alberta<sup>g</sup> |  |  |  | (Voordouw *et al.*, 2009) |  |
| *Mesotoga* sp. 3PWK154PWL11 | SC_PW.1 | 876,625 | 46.8 | JMRN01000000 | 21% |
| *Mesotoga* sp. 3PWM13N19 | SC_PW.2 | 1,886,634 | 45.8 | JMRM01000000 | 78% |
| *Mesotoga* sp. 4PWA21 | SC_PW.3 | 1,541,163 | 45.9 | JMQL01000000 | 34% |
| Oil sands tailings pond sediments near Fort McMurray, Alberta |  |  |  | (Tan *et al.*, 2015) |  |
| *Mesotoga* sp. NapDC | SC_NapDC | 1,885,291 | 45.8 | JNFM01000000 | 88% |
| *Mesotoga* sp. NapDC2 | SC_NapDC2 | 1,337,305 | 45.6 | JQSC01000000 | 53% |
| *Mesotoga* sp. NapDC3 | SC_NapDC3 | 1,828,922 | 45.6 | JWIP00000000 | 66% |
| Contaminated aquifer fluids from Colorado, USA |  |  |  | (Gieg *et al.*, 1999; Fowler *et al.*, 2012) |  |
| *Mesotoga* sp. TolDC<sup>h</sup> | SC_TOLDC | 2,257,992 | 46.1 | AYSI01000000 | 74% |

<sup>a</sup> Completeness of single cell genomes was calculated based on HMM hits to 119 single copy marker genes (see Supporting Material).
<sup>b</sup> Belongs geologically to the Glauconitic formation.
<sup>c</sup> These genomes were sequenced using IonTorrent PGM; all other genomes and single cells were sequenced using Illumina MiSeq
<sup>d</sup> Belongs geologically to the Upper Mannville Group – Cretaceous age.
<sup>e</sup> The *M. prima* genome was sequenced by Zhaxybayeva *et al.* 2012.
<sup>f</sup> The source was a sister-culture of the enrichment culture that yielded *M. prima* MesG1.Ag.4.2 (Nesbø *et al.*, 2006; 2012).
<sup>g</sup> Belongs geologically to the Western Canadian Sedimentary Basin.
<sup>h</sup> In addition to the SAG sequences, 39 *Mesotoga* fosmid clones prepared from the same culture were included in the assembly (http://hmp.ucalgary.ca/HMP/metagenomes/isolates.html; accessed October 2018).

The pan-genome of the *Mesotoga* isolate genomes was estimated to be 7,452,537 bp with an accessory genome of 5,664,475 bp; each genome contained a considerable amount of lineage-specific DNA (Fig. S1; see Supporting Information for additional details of the pan-genome and within-sample site diversity). In pairwise comparisons, the genomes shared on average 77% of their genes (Supporting Table S1).

### Phylogenetic analysis reveals three distinct *Mesotoga* lineages

The 16S rRNA genes of all 17 genomes had ≥99% identity to *M. prima*; phylogenetic trees revealed three distinct lineages (Fig. 1a). Genome networks based on core single nucleotide polymorphisms (SNPs) also had topologies consistent with the 16S rRNA gene phylogeny, with three distinct lineages being evident (Fig. 1b). Two lineages have a widespread geographical distribution: the World lineage (W; found in all regions represented) and the US lineage found in Baltimore Harbor and Colorado in the USA. The Alberta (A) lineage was observed in the Albertan samples only. Interestingly, *M. prima* has one 16S rRNA gene from the W lineage and one from the US lineage, suggesting one copy has been acquired laterally.

**Fig. 1.**
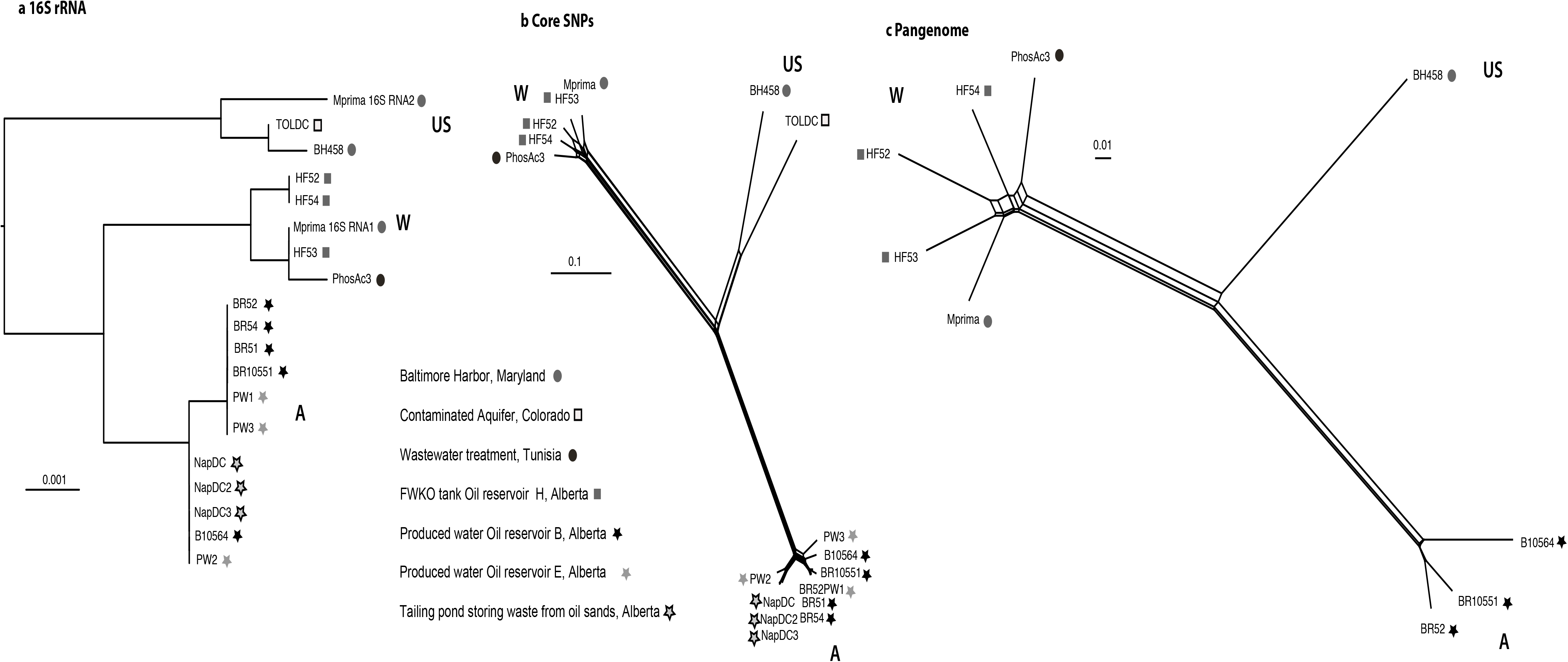
Phylogenetic relationships among *Mesotoga* genomes based on (a) 16SrRNA genes, (b) core SNPs and (c) presence/absence of shared 500-bp genomic fragments. The 16S rRNA maximum likelihood phylogeny was estimated using RAxML in Geneious v 10. For networks shown in (b) and (c), data were obtained using PanSeq (Laing *et al.*, 2010). Core SNPs in (b) were required to be present in 14 of 18 genomes (including SAGs), and genomic fragments were considered shared if they were at least 70% identical. The network in (c) was constructed using only genomes from isolates; shared fragments were required to be present in all 9 genomes and be at least 70% identical in nucleotide sequence. Networks were calculated in SplitsTree using NeighborNet algorithm (Huson and Bryant, 2006) from uncorrected distances. The isolates cluster into the same three lineages in (a), (b) and (c) and are named based on their geographical distribution. The World (W) lineage occurs in all regions represented. The US lineage is found in locations in the USA and the Alberta (A) lineage was observed in the Albertan samples only.

Very little reticulate evolution was observed among the three groups (Fig. 1b), and the A lineage in particular showed very little connection with the other two groups, suggesting that the three lineages have evolved independently for a relatively long time. In agreement with this, the ANI within groups ranged from 89.6-99.9%, while ANI between lineages ranged from 79.3-87.6% (Supporting Table S2). The same pattern was observed for the pangenome, with most lateral connections occurring within groups (Fig. 1c). Moreover, genomes from isolates of the same lineages share more genes in comparative analyses: average 86% within W and 92% within A (Supporting Table S1). Comparing genomes from different lineages, the US lineage had an intermediate position, sharing more genes with the A and W lineages: on average, genomes from A and W share 70% of genes, W and US share 76%, and A and US share 75% of their genes.

A high level of recombination was detected, with the majority (> 200) of recombination events involving genomes from the same lineage (Fig. S2). For the W and A lineages, respectively, the average recombination tract length was estimated to be 36,000 – 56,000 bp and 17,000–23,000 bp; the population mutation rate (θ) was estimated to be 0.022 and 0.013, and the population recombination rate (γ) to be 1.8 (range 1.5–2.2) and 1.5 (1.3–1.7). The resulting high γ/θ ratios of ~82–115 indicate high levels of recombination and are similar to estimates for *Thermotoga* spp. (Nesbø *et al.*, 2015).

Phylogenetic analysis identified 52 regions where recombination likely occurred between lineages: 39 regions showed evidence of recombination between *Mesotoga* sp. BH458 and the W lineage, eight regions suggested recombination between *Mesotoga* sp. BH458 and the A lineage, and only five regions showed possible recombination between A- and W-lineage genomes (Fig. 2). The regions with recombination involving the A lineage were short (range 230–530 bp) and the sequences more divergent, whereas several of the fragments involving the W lineage and *Mesotoga* sp. BH458 were > 5 kb (average 3000 bp, range 260–20,000). Multiple recombination events in the same locus will eventually result in shorter recombinant fragments being detected (see, e.g. (Mau *et al.*, 2006)). Taken together with the >10 kb length of the recombinant fragments detected in the within-lineage analysis, this difference in recombinant-fragment-length suggests that recombination events between the W lineage and *Mesotoga* sp. BH458 are more recent than those involving the A lineage. Very high levels of recombination were observed for a few genes. Among these is Theba_0319 in *M. prima*, the fourth most highly expressed gene (Supporting Table S3) that encodes the OmpB protein (Petrus *et al.*, 2012), a major component of the toga structure of Thermotogae.

**Fig. 2.**
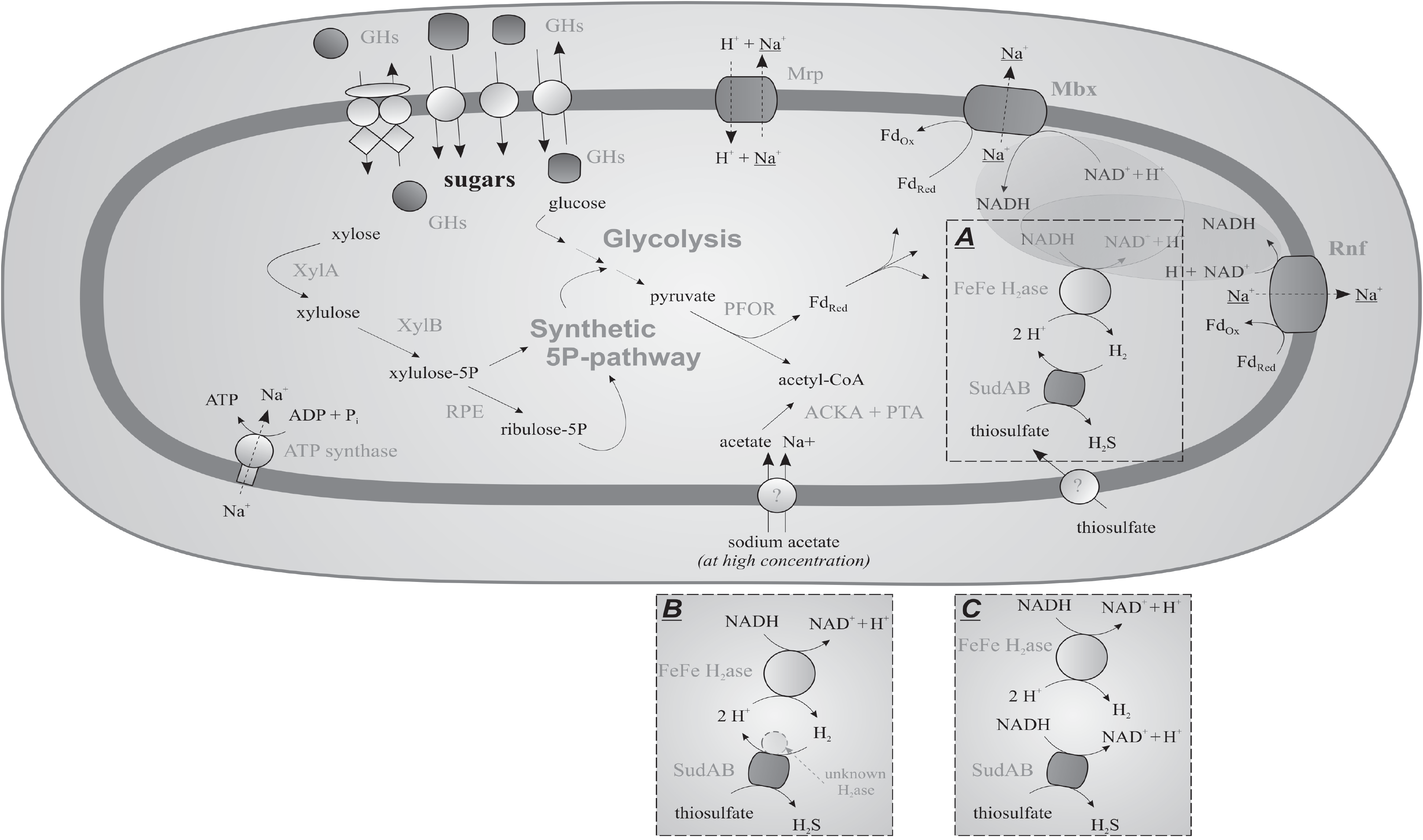
Visualization of recombination events detected among *Mesotoga* genomes from different lineages. The genomes are color-coded according to lineage (see text and Fig. 1) and arranged on the circumference of the circle: W lineage, blue; US lineage, orange; A lineage, green. Only isolate genomes were included in this analysis. A single representative genome (BR5.2) selected from the three highly similar genomes comprising the BR population (as described in Supporting Material) was included in the analysis. The recombination events with predicted donor and recipient are shown as lines connecting the two genomes; the locations of recombined regions, where line color reflects the donor lineage and the width of the line is proportional to the length of the recombinant region. The diagram was generated using Circos Version circos-0.69 (Krzywinski *et al.*, 2009).

### Comparison to metagenomes and phylogeographic patterns of the three *Mesotoga* lineages

We expanded the *Mesotoga* sequence dataset by searching IMG/M (in JGI) and SRA (in NCBI) databases for metagenomes containing *Mesotoga* spp. sequences. Fifteen metagenomes containing sequences closely related to the *Mesotoga* genomes investigated here were identified, arising from two environments already described (tailings pond and oil reservoir in Alberta), as well as oil reservoirs, contaminated sediments, wastewaters and hotspring sediments across the continental USA, and wastewaters in China (Table 2 and Supporting Information).

**Table 2.**
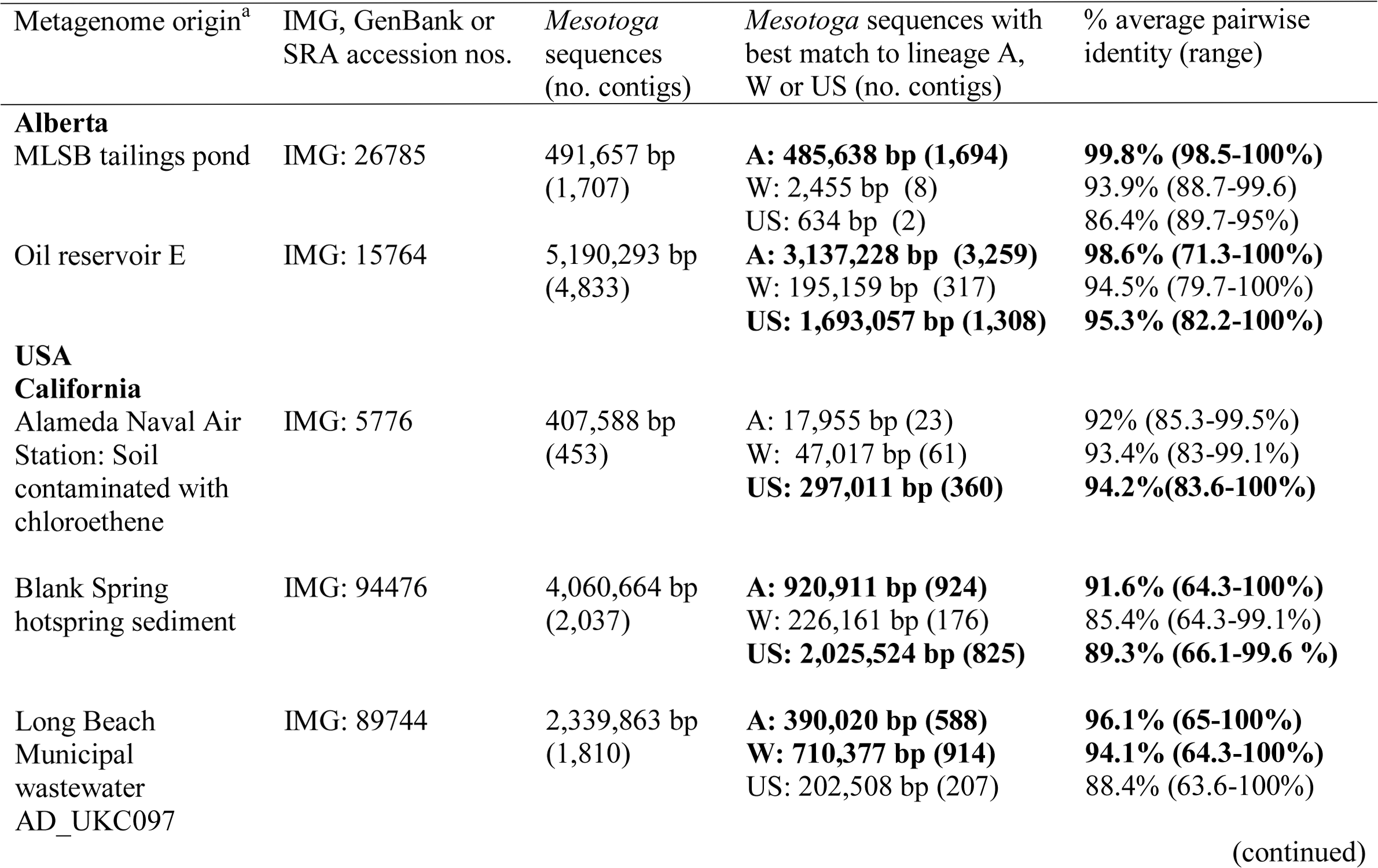
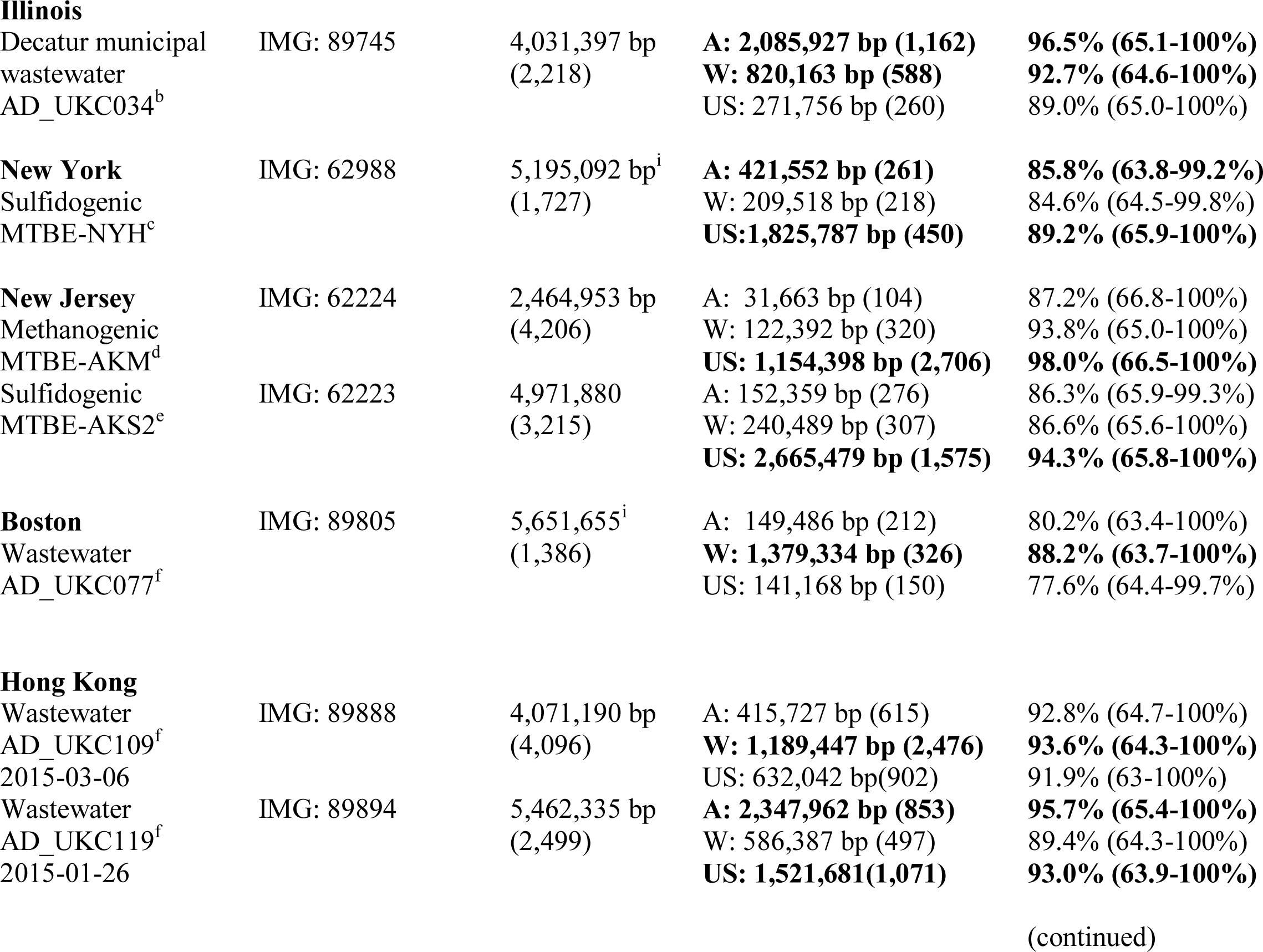
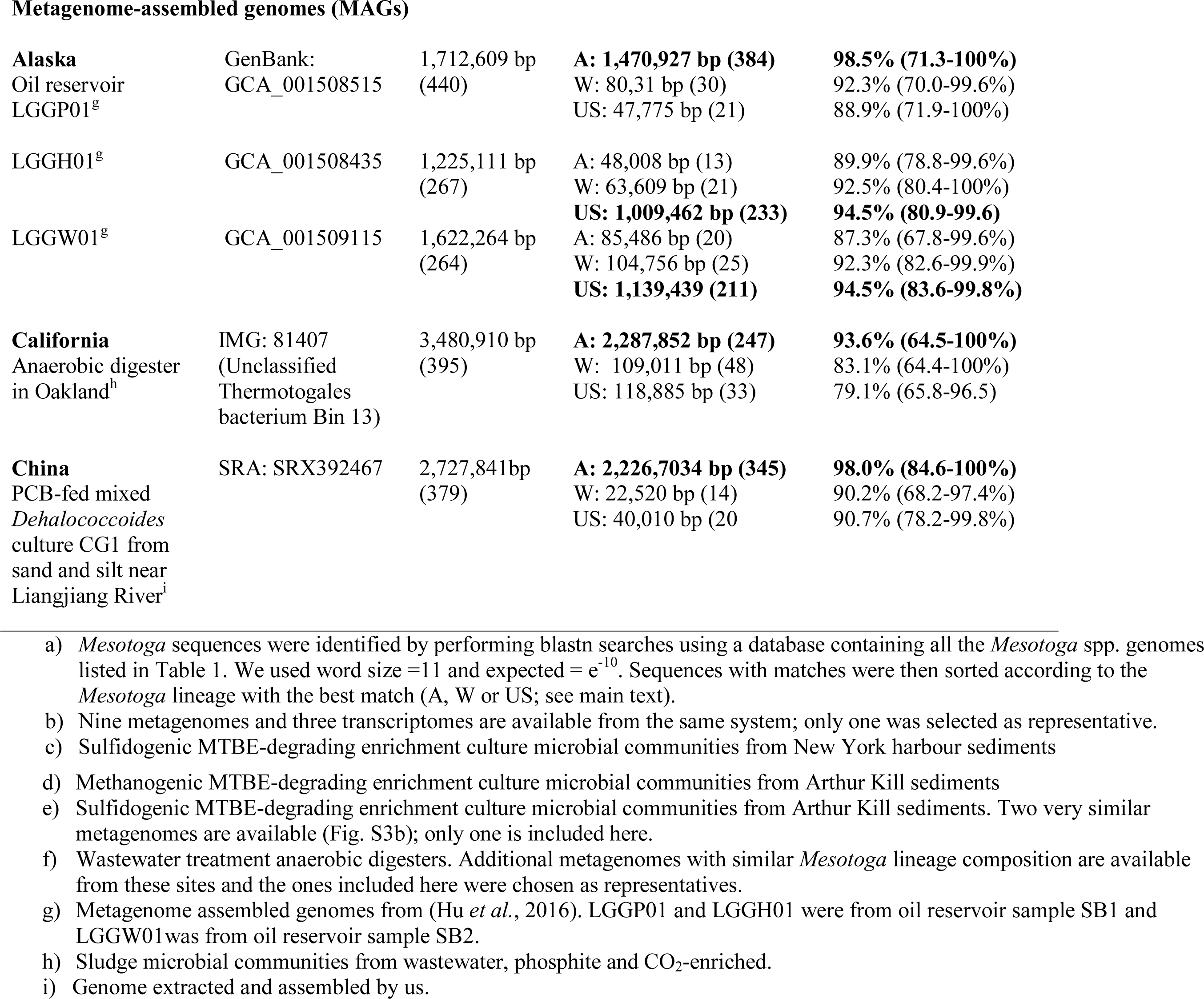
*Mesotoga* sequences recovered from publicly available metagenomes. For the sequences obtained from the IMG (JGI) database, only sequences classified as Thermotogae were downloaded. The predominant *Mesotoga* lineage extracted from each metagenome is shown in boldface. Metagenomes dominated by sequences similar to *Mesotoga infera* were not included.

#### Recent range expansion of the W-lineage

*Mesotoga* sequences with high similarity to the W lineage were identified using BLASTN searches in several wastewater treatment systems confirming its wide distribution in these environments (Table 2). A network including population genomes (PGs) of *Mesotoga* contigs (with > 90% sequence identity to W isolate genome) from three metagenomes dominated by W lineage sequences (Long Beach, Boston and Hong Kong, Table 2) revealed no geographical structuring.

#### Isolation by distance can explain the distribution of US genomes

The metagenome data expanded the observed distribution of the US lineage. As expected, metagenome IMG 15764 from Albertan oil reservoir E (the source of *Mesotoga* sp. SC_PW1-3) contained sequences with high identity to the A lineage. However, it also contained many sequences with high identity to the US lineage (Table 2), and sequence binning yielded two *Mesotoga* metagenome-assembled genomes (MAGs): one most similar to US-genomes (Fig. S3b) and one with a mix of sequences from the A lineage and US lineage (not shown).

The network of US-*Mesotoga* including PGs composed of contigs from metagenomes in Table 2 (with sequence identity > 80% to US-isolate genomes) revealed three groups (Fig. S3b) where PGs from New York and Blank Spring (California) form a cluster that does not contain any of the genomes sequenced in this study (Table 1). The clustering of remaining genomes correlates with both geography and environment type: the MAG assembled from oil reservoir E (Alberta), two MAGs from an Alaskan oil reservoir (Hu *et al.*, 2016), and the *Mesotoga* sequences from Alameda (California) clustered with SC_TOLDC from Colorado (western North America), while the *Mesotoga* sequences from New Jersey clustered with *Mesotoga* sp. BH458 from Baltimore Harbor (eastern North America). We therefore suggest that the divergence patterns seen for this lineage can be explained at least partly by an isolation-by-distance model.

#### Evolution of the A-lineage in isolation in North-American oil reservoirs

The metagenome sequences revealed that the A lineage is not restricted to Alberta, nor is it specific to oil reservoirs (Table 2), having substantial numbers of A-lineage sequences detected in wastewater metagenomes. For this lineage, MAGs were available from the same oil reservoir in Alaska where we observed the US-lineage (Hu *et al.*, 2016), an anaerobic wastewater digester in Oakland (California), and one, assembled by us, from a PCB-fed culture inoculated with sediments from Liangjiang River, China (Wang and He, 2013). Network analysis revealed that the genome from the Alaskan oil reservoir is most similar to those from the Albertan oil reservoir B, whereas the genomes from China and California show high similarity (> 99%) to each other and to *Mesotoga* sp. SC_NapDC from a northern Albertan oil sands tailings pond (Fig. S3c).

### Distinct metabolism in mesophilic Thermotogae

We also examined the newly available genomes for metabolic insights, which may be linked to *Mesotoga*’s lower growth temperatures and may influence the role(s) *Mesotoga* play in their environments.

#### *Mesotoga*-specific genes

Comparison of the *Mesotoga* isolate genomes to other Thermotogae genomes in IMG revealed 200 *M. prima* genes found in all *Mesotoga* genomes (including the more distantly related *Mesotoga infera* not included in the phylogenomic analyses), but in no other Thermotogae genomes. The majority of these genes were hypothetical proteins (N=119, Supporting Table S3). When *Mesotoga*-specific genes with a predicted function were classified according to Clusters of Orthologous Groups (COG) categories, the largest category was ‘Amino Acid metabolism and transport’ with 11 genes, most of which were dipeptidases (COG4690, N=6).

#### Mesotoga-specific genes related to O_2_ exposure

Several *Mesotoga*-specific genes are predicted to be involved in oxygen radical defense (Supporting Table S4). One of the most highly conserved genes across all the *Mesotoga* genomes (Theba_1553; average pairwise identity 96.3%) shows similarity to peroxiredoxin and alkyl hydroperoxide reductase domain-encoding genes. Moreover, a catalase gene (Theba_0075) is found in all isolate genomes except those from oil reservoir H.

#### Reducing equivalents and thiosulfate reduction

*Mesotoga*’s core metabolism differs from that of other characterized Thermotogae. While growth of most Thermotogae is stimulated by, but not dependent upon, the presence of thiosulfate, sulfur, or other reduced sulfur compounds in laboratory medium (Ravot *et al.*, 1995; Boileau *et al.*, 2016), reduction of sulfur compounds appears to be essential for growth of *Mesotoga* in pure culture (Hania *et al.*, 2011; 2013; Fadhlaoui *et al.*, 2017). The first description of *M. prima* (Nesbø *et al.*, 2012) reported that growth was only slightly stimulated by the presence of thiosulfate or sulfur. However, here we observed growth of this isolate only in the presence of sulfur or thiosulfate (Supporting Table S5 and Table S6), confirming that this is a general trait of *Mesotoga* spp. Additionally, while other Thermotogae produce H_2_ (and H_2_S if grown with partially reduced sulfur compounds), *Mesotoga* spp. produce large amounts of H_2_S and no or little H_2_ (Supporting Table S5).

To reconcile these observations with genomic data, transcriptome analysis was performed using a culture of *M. prima* grown with 0.5% yeast extract, xylose and thiosulfate. RNAseq analysis revealed high expression of Theba_0443 (RPKM of 3650; Supporting Tables S3 and S4) encoding a Fe-hydrogenase homologous to the one used by *Kosmotoga olearia* (Kole_0172). Hydrogenases are indeed essential in Thermotogae for recycling of ferredoxins (Schut *et al.*, 2013); therefore, finding the same hydrogenase to be highly expressed in *M. prima* and *K. olearia*, and conserved in all *Mesotoga* genomes investigated here, suggests that *Mesotoga* possesses a mechanism relying on oxidized sulfur compounds, efficiently converting all intracellularly produced H_2_ to H_2_S. Notably, there was no change in the culture headspace gas H_2_:N_2_ ratio after incubating *Mesotoga* spp. in a 1:9 H_2_:N_2_ atmosphere for > 5 months (Supporting Table S5), suggesting that *Mesotoga* neither produces nor takes up externally supplied H_2_.

No homologs of characterized thiosulfate reductases were identified, although the *Mesotoga* genomes carry homologs (Theba_0076; Theba_0077 in *M. prima*) of an archaeal intracellular ferredoxin:NADP oxidoreductase (SudAB; (Hagen *et al.*, 2000)) capable of acting as a sulfide dehydrogenase in the presence of elemental sulfur or polysulfide (Fig. 3). Both genes were transcribed at moderate levels in *M. prima* grown with thiosulfate (RPKM 341 and 243, respectively), whereas the *K. olearia* homologs (Kole_1827, Kole_1828) were highly expressed under similar conditions (RPKM > 1000, (Pollo *et al.*, 2017). SudAB complexes, however, are not known to be involved in thiosulfate reduction. This is probably due to an unfavorable *E*°=82 mV for the reaction when NADH acts as electron donor: *E*°` [S_2_O_3_ ^2-^/ HS^-^ + SO_3_ ^2-^] = -402 mV and *E*°` [NAD+/ NADH] = -320 mV. The *E*°` of [Fd_Ox_ / Fd_Red_] is similarly high at -390 mV. Comparable endergonic reactions are catalyzed by the *Salmonella enterica* thiosulfate reductase (Phs) by utilizing proton-motive force (Stoffels *et al.*, 2012). However, the cytoplasmic SudAB complex cannot couple proton-motive force and reduction of an external electron acceptor. Thus, neither NADH nor Fd_Red_ can function as electron donors for thiosulfate reduction by *M. prima*. Instead molecular H_2_ with *E*°` [2H^+^ / H_2_] = -410 mV appears to be a thermodynamically preferable electron donor for thiosulfate reduction. The only hydrogenase present in the *M. prima* genome is the highly expressed FeFe-hydrogenase (Theba_0443), which usually is involved in Fd-dependent H_2_ production (Vignais and Billoud, 2007). However, a cluster of five highly transcribed genes (Theba_0461 – 0465, RPKM 1203-3697, Supporting Table S4) encodes proteins homologous to all subunits of the NADP-reducing hydrogenase Hnd of *Desulfovibrio fructosovorans* (Nouailler *et al.*, 2006) except the catalytic subunit (HndD). These proteins may work together with Theba_0443 to form a FeFe-hydrogenase complex (Fig. 3). We hypothesize that this complex is involved in intracellular synthesis of molecular hydrogen for thiosulfate reduction by SudAB coupled to NADH oxidation (formed by Mbx and/or Rnf complexes, see below and Fig. 3). The Hnd genes have homologs in other Thermotogae, however, similar genomic context is observed only in genomes of other *Mesotoga* and *Kosmotoga* spp. (Supporting Table S7).

**Fig. 3.**
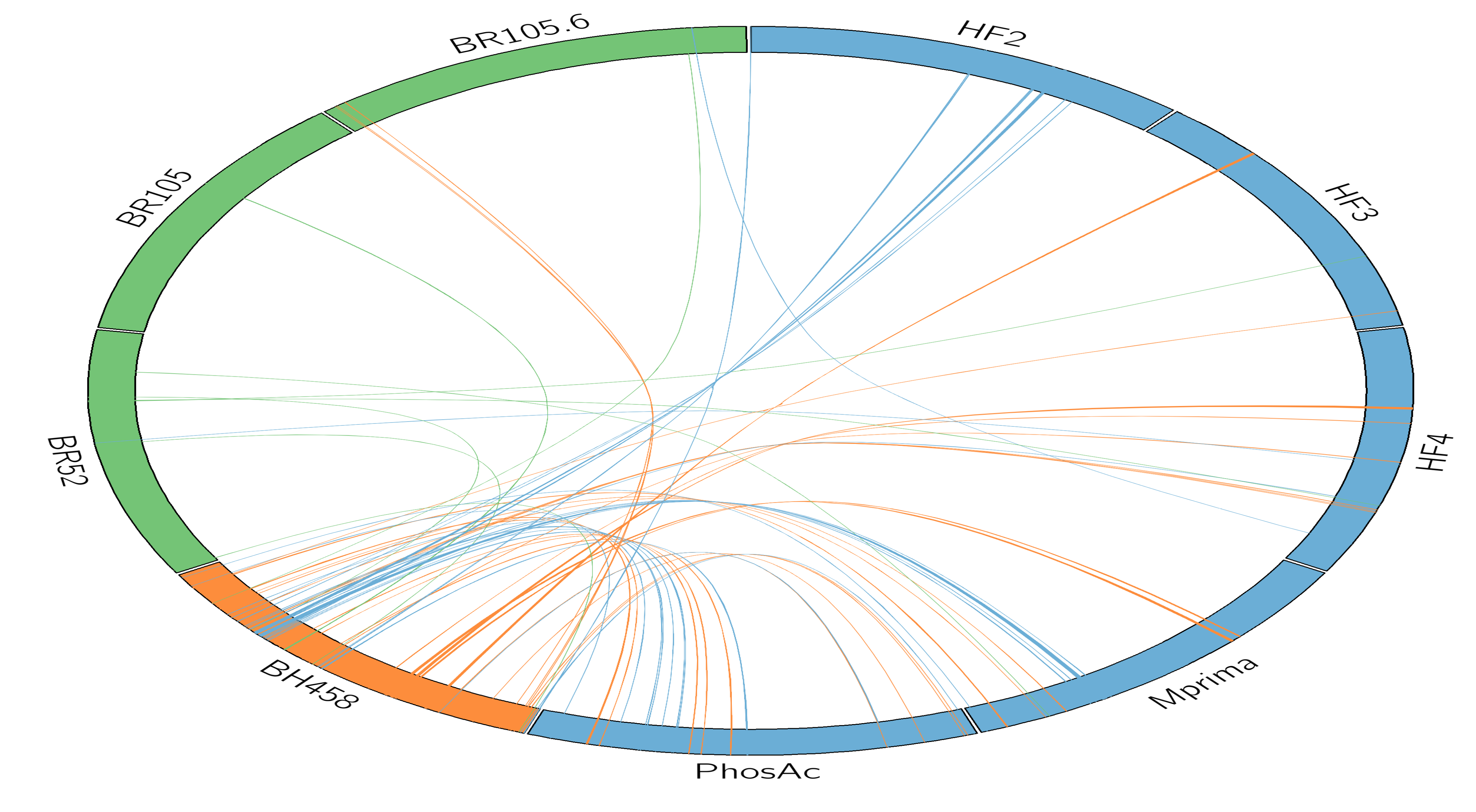
Model of energy generation pathway in *Mesotoga prima* during growth on xylose and thiosulfate. Glucose and xylose poly- and oligosaccharides are hydrolyzed by various intracellular and interstitial glycosidases (GHs). Glucose oxidation occurs via the glycolytic Embden-Meyerhof-Parnas pathway, whereas xylose is utilized via xylose isomerase (*XylA*, Theba_1394), xylulose kinase (*XylB*, Theba_1395, Theba_2230, Theba 2429, Theba 2518, Theba 2544, Theba 2588), ribulose phosphate 3-epimerase (Theba_0639) and enzymes of the pentose-phosphate pathway. Specifically, xylose isomerase converts D-xylose to D-xylulose, which is phosphorylated by the set of xylulose kinases to D-xylulose 5-phosphate, and further to ribulose 5-phosphate by the ribulose-phosphate 3-epimerase. Both xylulose 5-phosphate and ribulose 5-phosphate produced by this pathway are common metabolic intermediates in the pentose phosphate pathway. Xylose isomerase (Theba_1394) was among the most highly transcribed genes during cultivation of *M. prima* on xylose and thiosulfate (Supporting Table S3). Acetyl-CoA formation occurs by means of pyruvate-ferredoxin oxidoreductase (PFOR, Theba_1954). In the possible case of growth on acetate, its activation occurs by means of acetate kinase (ACKA, Theba_0428) and phosphotransacetylase (PTA, Theba_0782), acting in reverse. The model includes gene products hypothesized to be involved in thiosulfate reduction. <u>Na^+^</u> refers to Na^+^ ions involved in generating sodium motive force. A: The FeFe hydrogenase (Theba_0443 and Theba_0461 – 0465) reduces NADH to form H_2_, which is used as an electron donor for thiosulfate reduction catalyzed by SudAB (Theba_0076, Theba_0077). Mbx (Theba_1796-1808) and/or Rnf (Theba_1343-1348) complexes provide additional NADH along with the oxidation of excessive reducing equivalents (Fd_red_) and generation of a sodium motive force. B and C: other possible scenarios of H_2_ oxidation and thiosulfate reduction.

*Mesotoga* cells require enzymes that re-oxidize Fd_red_ formed during sugar oxidation. This might be carried out by either the NADP:ferredoxin oxidoreductase complex (Mbx; Theba_1796-1808 in *M. prima,* (Schut *et al.*, 2013)) or the Rnf ion-motive electron transport complex (Theba_1343-1348; (Müller *et al.*, 2008). Conserved motifs (Mulkidjanian *et al.*, 2008) suggested a Na^+^-translocating F-type ATP synthase operating in *M. prima.* As a consequence, both Mbx and Rnf complexes are predicted to export Na^+^ generating sodium-motive force instead of proton-motive force. Genes encoding Mbx and Rnf show low and moderate expression (RPKM 37-88 and 236-478, respectively) during growth on thiosulfate, and the expression values suggests that Rnf is the main complex involved.

#### Acetate and xylose utilization

Growth on acetate was reported for *Mesotoga* PhosAc3 (Hania *et al.*, 2015), and we observed weak stimulation of growth of its close relative *M. prima* by acetate (day 5-10 in Supporting Fig. S4 and Table S6). (Nobu *et al.*, 2015) suggested that Ca. “*Mesotoga acetoxidans*”, a MAG closely related to *M. infera,* oxidizes acetate by using a novel pathway even though the genes comprising the pathway are conserved in all Thermotogae genomes. Yet, this phenotype is uncommon among Thermotogae and has been reported only for *Pseudothermotoga lettingae* (Balk *et al.*, 2002). Instead, many Thermotogae are inhibited by acetate, including one of *Mesotoga*’s closest relatives, *K. olearia* (Dipippo *et al.*, 2009). Our search for *Mesotoga*-specific genes that may be responsible for their observed growth on acetate revealed a candidate gene encoding a bacterial homodimeric pyruvate:ferredoxin oxidoreductase (PFOR; Theba_1954), with close homologs only found in *Kosmotoga pacifica* (Jiang *et al.*, 2017) and *Mesoaciditoga lauensis* (Reysenbach *et al.*, 2013). Unfortunately, the description of these two species did not investigate growth on acetate. The *pfor* gene is distantly related to the archaeal multi-subunit-type used by other Thermotogae (Ragsdale, 2003) and almost all its close homologs fall within the *Clostridia* (Supporting Fig. S4). Genes having 97-99% identity to *pfor* from *M. infera*, and 83-85% identity to the *M. prima* homolog, were found in both the metagenome and metatranscriptome published by (Nobu *et al.*, 2015) (locus tag JGI12104J13512_10052834 and JGI11944J13513_10066464) but were not included in their model. We propose that PFOR may work with the acetate kinase (Theba_0428 in *M. prima*) and phosphotransacetylase (Theba_0782 in *M. prima*) found in all Thermotogae to enable *Mesotoga* to grow on acetate. At high extracellular acetate concentrations we suggest that PFOR shifts the balance favoring the production of pyruvate from acetyl-CoA (i.e. serves as an acetate switch (Wolfe, 2005)).

*M. prima* grows optimally on xylose, a sugar fermented by many Thermotogae (Bhandari and Gupta, 2014). The D-xylose utilization pathway is similar to that observed in Firmicutes (Gu *et al.*, 2010) (Fig. 3). Several possible xylulose kinase genes were found co-localized with genes encoding xylosidases, sugar transporters, and kinases, suggesting their synergetic activities in xylan hydrolysis, xylose import, and utilization.

## Discussion

### *Mesotoga* have conserved core genomes and diverse pangenomes

The comparative analysis of the *Mesotoga* genomes revealed higher levels of diversity in genome content than observed in the hyperthermophilic Thermotogae. Whereas *Thermotoga* spp. share > 90% of their genes in pairwise comparisons (Nesbø *et al.*, 2015), *Mesotoga* genomes from the same lineage share on average 86% - 92% of their genes. (Nesbø *et al.*, 2015) suggested that high levels of recombination may be partly responsible for homogenizing *Thermotoga* spp. genomes. However, since we observed similar high levels of recombination within the *Mesotoga* W and A lineages, additional forces must be responsible for the larger proportion of variable accessory genes. Perhaps more cryptic niches are available in low- versus high-temperature subsurface environments (McInerney *et al.*, 2017), or *Mesotoga* may have larger effective population sizes than the hyperthermophiles (Andreani *et al.*, 2017).

Comparing the nucleotide divergence within the core genomes revealed ‘species’ level divergence between the three lineages detected (ANI < 87%), while ANI within the A and W lineage was very high at 98.5% and 97.5%, respectively. In comparison, the ANI among the *Thermotoga* genomes investigated by (Nesbø *et al.*, 2015) was 95.3%. Thus *Mesotoga* spp., particularly those from the W-lineage, appear to have more conserved core genomes and more diverse pangenomes than their hyperthermophilic relatives.

### Three *Mesotoga* lineages with distinct phylogeographies: isolation by distance, range expansion, and burial with isolation

The networks calculated for both the core and the pangenome gave the same overall topology as that observed in the 16S rRNA tree with three distinct groups. The low level of recombination observed among these three groups suggests they have evolved independently for a relatively long time. The observation of several recent recombination events between the W and US lineages, which currently co-exist in at least one location (i.e., Baltimore Harbor), demonstrates that recombination between lineages is possible. We therefore suggest that the three *Mesotoga* lineages have evolved independently due to geographical, not genetic, isolation. This is contrary to the patterns of geneflow observed in *Thermotoga* spp. genomes, where environment type was more important than geographic separation in determining level of geneflow (Nesbø *et al.*, 2015). Although it may seem counterintuitive that mesophilic *Mesotoga* would be more affected by geographical separation than hyperthermophilic *Thermotoga*, this may be a consequence of their anaerobic metabolism. (Chakraborty *et al.*, 2018) showed that bacteria are dispersed out of deep hot subsurface oil reservoirs and into the ocean through hydrocarbon seeps, and this might serve as a major route of migration between these environments. Moreover, temperature gradients associated with hydrothermal systems are often very sharp (Dick *et al.*, 2013), and hyperthermophilic *Thermotoga* cells will therefore quickly become inactive if they enter cold aerobic ocean water (Fig. S6). Mesophilic *Mesotoga* cells will, however, more likely enter oxygenated environments having a suitable temperature before they reach a new optimal anaerobic site and therefore may more often succumb to oxygen exposure, limiting viable dispersal and gene exchange (Fig. S6). In support of this, many *Mesotoga*-specific genes appear to be involved in O_2_ or H_2_O_2_ detoxification.

Within the three lineages we see patterns consistent with different phylogeographic histories. Comparing the isolate genomes to *Mesotoga* sequences in metagenomes, the US-lineage shows patterns consistent with isolation by distance. Moreover, the US-lineage has an intermediate position between the A- and W-lineages when considering ANI, gene content, and recombination, which may be due to this lineage co-existing with both W and A genomes (e.g. Baltimore Harbor, Oil field E).

Members of the widespread W-lineage show high identity in their core genomes, large pan-genomes, and no indication of geographical structuring, indicative of a recent range expansion (Choudoir *et al.*, 2017). To date, W lineage *Mesotoga* have been detected only at sites heavily influenced by human activities (e.g., drilling, contamination), suggesting an anthropogenic role in their dispersal and possibly selective pressure on these genomes. Interestingly, one of the W-lineage-specific genes (Theba_0620, Supplemental material) is involved in synthesis of poly-gamma glutamate, which has been implicated in survival under harsh conditions and may have contributed to the wide distribution of this lineage.

The A lineage is more isolated from the other lineages (Fig. 1 and 2), which might suggest that this clade evolved in isolation since the formation of oil reservoir sediments in Alberta 55–120 Ma (Schaefer, 2005; Selby, 2005; Head *et al.*, 2014). The high similarity of the MAGs from the Alaskan oil field to the Albertan genomes and MAGs from the A and US lineages (Fig. S3) could be due to these oil reservoir sediments being laid down around the same time (~100 Ma (Hu *et al.*, 2016). However, the position of these MAGs in the genome networks could also be explained by these oil reservoirs being colonized by the same subsurface population, as suggested for *Thermotoga* spp. (Nesbø *et al.*, 2015). Additional oil reservoir genomes are needed to resolve this question and also to determine if the A-lineage is indeed indigenous to oil reservoirs.

Also this lineage has likely experienced recent dispersal events due to human activities: MAGs from a polluted river bank in Liangjiang, China (Wang and He, 2013) and waste water from Oakland (California) showed very high identity to *Mesotoga* sp. SC_NapDC from a northern Albertan oil sands tailings pond. In fact, these genomes show the second highest level of pairwise identity among the A lineage genomes (Fig. S3d), suggesting recent dispersal, possibly due to human activities in the last decades.

### Distinct metabolism in mesophilic Thermotogae

The *Mesotoga* genomes and transcriptome also elucidated the genetic background for their distinct energy metabolism compared to thermophilic Thermotogae bacteria, i.e. the strict need for sulfur or thiosulfate and no or little H_2_ production, but rather H_2_S production unless in co-culture with a sulfate reducer (Fadhlaoui *et al.*, 2017). (Fadhlaoui *et al.*, 2017) suggested that *Mesotoga’*s inability to ferment sugars is mainly due to its lack of a bifurcating hydrogenase. However, *K. olearia* also lacks this enzyme and ferments pyruvate, producing large amounts of hydrogen using the homolog of *M. prima*’s only Fe-hydrogenase (Pollo *et al.*, 2017). In the model in Fig. 3 panel A, we therefore instead suggest this is accomplished by utilizing a bifurcated hydrogenase, SudAB, Mbx and Rnf.

The model shown in Fig. 3 panel A accounts for the observed dependence of *M. prima* on sulfur or thiosulfate for growth, the lack of H_2_ production, and involves proteins previously implicated in hydrogen and sulfur metabolism. Importantly, however, currently there are no known enzymes that couple H_2_ oxidation and thiosulfate/sulfur reduction. It is therefore possible that *M. prima* SudAB uses NADH as the electron donor and is much more effective than the hydrogenase which results in almost no H_2_ as growth product (Fig. 3 panel C).

Alternatively, thiosulfate reduction coupled to H_2_ oxidation (i.e., the postulated role of SudAB; Fig. 3 panel A) may be performed solely by one of the highly-transcribed hypothetical *Mesotoga* proteins with no match to genes in *Kosmotoga* and other Thermotogae, or in combination with SudAB (Fig. 3 panel B). Several candidate genes listed in Supporting Table S4 encode proteins with unknown functions. Functional studies of these genes, as well as the gene products shown in Fig. 3, are needed to assess their role, if any, in thiosulfate reduction. Additional genes that may be involved in thiosulfate uptake and electron transfer are also discussed in Supporting Information. Interestingly, PhosAc3 and *M*. *infera* were found to reduce only elemental sulfur (Hania *et al.*, 2011; 2013) whereas the strains isolated by us also reduce thiosulfate. These differences may reflect selection during isolation; all the isolates obtained in the current study were from enrichment cultures containing thiosulfate, whereas PhosAc3 and *M. infera* were enriched with sulfur. This suggests that the sulfur-compound-preference may be a variable and flexible phenotype in *Mesotoga* populations.

We also observed gene content differences that probably are directly linked to *Mesotoga*’s lower growth temperature. The higher abundance of genes associated with oxygen radical defense may be linked to the lower growth temperatures of *Mesotoga* versus thermophilic Thermotogae. O_2_ solubility in water is greater and free radicals are stabilized at low temperatures, and organisms living at low temperatures are therefore exposed to higher concentrations of reactive oxygen species (Piette *et al.*, 2010). It should be noted that the transcriptome of *M. prima* grown anaerobically revealed that two of the genes possibly involved in O_2_ or H_2_O_2_ defense (Theba_0075, Catalase and Theba_2399, Rubrerythrin) were highly expressed (top 5% of expressed genes; Supporting Table S3 and S4), suggesting that these genes may have additional or alternative functions under anaerobic conditions. Further investigation is needed to clarify the transcriptional responses of these genes and identify the targets of their enzymes.

## Conclusion

Our genomic analysis suggests that the lower growth temperature of *Mesotoga* spp. compared to the hyperthermophilic *Thermotoga* has likely influenced *Mesotoga* phylogeography, with geographic separation historically having a greater influence than genetic separation, possibly due to the damaging effects of oxygen exposure during dispersal (Fig. S6). Whether this is a general feature of strictly anaerobic organisms remains to be resolved. There is also some indication of possible ecotype differentiation among the *Mesotoga* lineages, with the US lineage being common in communities degrading aromatic pollutants (PCB, toluene) and the A lineage in hydrocarbon-impacted sites. However, for both of these lineages, inspection of metagenomes revealed they are not restricted to these environments. The analysis including data from metagenomes also suggests that anthropogenic activities have expanded *Mesotoga*’s habitats and also enhanced its dispersal capabilities (Fig. S6), with inferred recent long-distance dispersal events involving anthropogenic environments and/or activities.

The ecological role of *Mesotoga in situ* may differ from their thermophilic relatives. For instance, hydrogen-producing *Thermotoga* spp. have been shown to grow in syntrophy with hydrogenotrophic methanogens (e.g., (Johnson *et al.*, 2005)) but this is likely not the case for *Mesotoga* that produce only trace amounts or no detectable extracellular H_2_. Supporting this proposal, we were unable to establish co-cultures of *M. prima* and a hydrogenotrophic methanogen (not shown). Instead (Fadhlaoui *et al.*, 2017) showed that *Mesotoga* spp. prefer to grow in syntrophy with hydrogenotrophic sulfate-reducing bacteria. This, together with the ability to both produce and consume acetate, suggests that *Mesotoga* will assume different environmental roles than their thermophilic relatives, for instance by supporting the growth of sulfate reducers rather than methanogens. An interesting question is whether they also grow syntrophically with other common hydrogenotrophic organisms in their niches, such as organohalide-respiring *Dehalococcoides* (e.g. (Fagervold *et al.*, 2007)). Finally, the large amounts of H_2_S produced by *Mesotoga* could have detrimental effects on oil reservoirs, production facilities, and pipelines where *Mesotoga* is commonly found. Monitoring the presence of *Mesotoga* spp. in addition to the more commonly targeted sulfate reducers in these industrial environments (Lee *et al.*, 1995) may be informative and valuable.

## Experimental procedures

### Sources of genome sequences

Nine *Mesotoga* strains (BR, HF and BH designations) were isolated from oil reservoirs and anaerobic sediments in Canada and the USA (Table 1). All nine available isolates were selected for genome sequencing. In addition, seven single cells were physically selected from oil field fluids or oil sands enrichment cultures from Canada or a contaminated aquifer in the USA (PW, NAPDC and TOLDC designations, respectively) and amplified by PCR to produce SAGs. Detailed descriptions of isolation procedures, DNA extraction, genome assembly and annotation are provided in Supporting Information.

To augment the strain genomes, 15 publicly available metagenomes containing large numbers of *Mesotoga* spp. sequences were identified using blastn searches of IMG (JGI; accessed February 2017) and SRA (NCBI; accessed December 2016) using *rpoB* from *M. prima* as a probe and expected (exp.) set to < e^-50^. For additional details on search parameters and information on assembly of draft genomes from metagenomic sequences or contigs see Supporting Information.

### Genome content and genome alignments

Shared genes and genome specific genes were identified in IMG Version 4 (Markowitz *et al.*, 2014) using translated proteins and 70% identity cut-off and exp. < e^-10^, whereas 30% sequence identity cut-off and exp. < e^-5^ were used to identify lineage-specific genes and for comparing *Mesotoga* genomes to other Thermotogae genomes.

Pan-genome calculations were performed in Panseq (Laing *et al.*, 2010) using a fragment size of 500 bp and 70% identity cutoff, and exp. < e^-10^. The data matrices of shared core SNPs and 500-bp fragments were converted into uncorrected distances and visualized in SplitsTree 4 (Huson and Bryant, 2006) using NeighborNet clustering.

Whole genome alignments were carried out in MAUVE version 2.3.1 (Darling *et al.*, 2010) using automatically calculated seed weights and minimum Locally Collinear Blocks (LCB) scores. LCB positions with gaps were removed and the edited LCB were concatenated in Geneious v.10 (www.geneious.com). Average nucleotide identities (ANI) were calculated at http://enve-omics.ce.gatech.edu/ani/ (Goris *et al.*, 2007).

### Recombination detection

The relative rate of recombination to mutation within lineages, as well as the average recombination tract length, were assessed using the LDhat package (McVean *et al.*, 2002; Jolley *et al.*, 2004) as described by (Nesbø *et al.*, 2015) on concatenated alignments (including LCB > 10,000 bp) of genomes from the W and the A lineage separately. Recombinant fragments between lineages were detected using LikeWind Version 1.0 (Archibald and Roger, 2002) on the concatenated MAUVE alignment (above), using a sliding window of 1000 bp with 100-bp increments.

### RNAseq analysis

RNA isolation from a culture of *M. prima* (grown at 45°C for 73 h in 0.5% yeast extract, M thiosulfate and 0.5% xylose) and subsequent sequencing as one of five barcoded libraries were performed as described by (Pollo *et al.*, 2017). RNAseq analysis was carried out in CLC Genomics Workbench version 7.0.4 as described by (Pollo *et al.*, 2017). The transcriptome has been submitted to GenBank’s SRA archive with accession number PRJNA495810.

### H_2_ and H_2_S measurements

Standard gas chromatographic analysis of culture headspace gas was performed using an Agilent CP4900 Micro Gas Chromatograph to detect H_2_ production by the cultures, as described in Supporting Information. Dissolved sulfide concentrations were measured using a VACUettes^®^ Visual High Range Kit (Chemetrics), following the manufacturer’s recommendations.

## Acknowledgements

This work was supported by a Research Council of Norway award (project no. 180444/V40) to C.L.N. and by a Genome Canada grant (Hydrocarbon Metagenomics Project) to J.F. The work of IVK was supported by the Russian Science Foundation grant # 18-44-04024. We thank Dr. Alexander Lebedinsky for constructive criticism and helpful suggestions.

